# *C9orf72* gene networks in the human brain correlate with cortical thickness in C9-FTD and implicate vulnerable cell types

**DOI:** 10.1101/2023.07.17.549377

**Authors:** Iris J. Broce, Daniel W. Sirkis, Ryan M. Nillo, Luke W. Bonham, Suzee E. Lee, Bruce Miller, Patricia Castruita, Virginia E. Sturm, Leo S. Sugrue, Rahul S. Desikan, Jennifer S. Yokoyama

## Abstract

**Introduction:** A hexanucleotide repeat expansion (HRE) intronic to chromosome 9 open reading frame 72 (*C9orf72*) is recognized as the most common genetic cause of amyotrophic lateral sclerosis (ALS), frontotemporal dementia (FTD), and ALS-FTD. Identifying genes that show similar regional co-expression patterns to *C9orf72* may help identify novel gene targets and biological mechanisms that mediate selective vulnerability to ALS and FTD pathogenesis.

**Methods:** We leveraged mRNA expression data in healthy brain from the Allen Human Brain Atlas to evaluate *C9orf72* co-expression patterns. To do this, we correlated average *C9orf72* expression values in 51 regions across different anatomical divisions (cortex, subcortex, cerebellum) with average gene expression values for 15,633 protein-coding genes, including 50 genes known to be associated with ALS, FTD, or ALS-FTD. We then evaluated whether the identified *C9orf72* co-expressed genes correlated with patterns of cortical thickness in symptomatic *C9orf72* pathogenic HRE carriers (n=19). Lastly, we explored whether genes with significant *C9orf72* radiogenomic correlations (i.e., ‘*C9orf72* gene network’) were enriched in specific cell populations in the brain and enriched for specific biological and molecular pathways.

**Results:** A total of 1,748 genes showed an anatomical distribution of gene expression in the brain similar to *C9orf72* and significantly correlated with patterns of cortical thickness in *C9orf72* HRE carriers. This *C9orf72* gene network was differentially expressed in cell populations previously implicated in ALS and FTD, including layer 5b cells, cholinergic motor neurons in the spinal cord, and medium spiny neurons of the striatum, and was enriched for biological and molecular pathways associated with multiple neurotransmitter systems, protein ubiquitination, autophagy, and MAPK signaling, among others.

**Conclusions:** Considered together, we identified a network of *C9orf72*-associated genes that may influence selective regional and cell-type-specific vulnerabilities in ALS/FTD.

## Introduction

Frontotemporal dementia (FTD) and amyotrophic lateral sclerosis (ALS) are neurodegenerative disorders. ALS and FTD show overlapping clinical, genetic, and neuropathological features. FTD is the most common form of dementia diagnosed in people younger than 65 years old and is characterized by changes in social behavior and/or language abilities due to neurodegeneration of the frontal and temporal lobes. Depending on the signs and symptoms, FTD patients are classified into one of three different syndromes: behavioral variant FTD (bvFTD) or one of two forms of primary progressive aphasias (PPA), including non-fluent variant PPA (nfvPPA) and semantic variant PPA (svPPA). ALS is the most common form of adult-onset motor neuron disease (MND) and is characterized by progressive degeneration of both upper motor neurons of the motor cortex and lower motor neurons of the brainstem and spinal cord at disease onset. Although motor neuron damage predominates in ALS, other neuronal populations including within frontal, temporal, and parietal cortical circuits, the basal ganglia, and dorsal root ganglia are also involved in some patients (Bang et al., 2015; Rabinovici & Miller, 2010; Rosness et al., 2008; van Vliet et al., 2013). Although the clinical phenotypes of FTD and ALS can be heterogeneous, about 15% of people with bvFTD, 11% of patients with nfvPPA, and 19% of patients with svPPA may eventually develop motor symptoms consistent with ALS (Rascovsky et al., 2011; Vinceti et al., 2019). Similarly, about 50% of ALS patients develop cognitive and behavioral impairment, with 13% meeting diagnostic criteria for bvFTD (Ljubenkov & Miller, 2016). This clinical overlap may, at least in part, be due to shared neuropathology due to aggregation of TDP-43, which drives MND and around half of frontotemporal lobar degeneration (FTLD) pathology. After decades of research, it is now recognized that a pathogenic hexanucleotide repeat expansion (HRE) intronic to chromosome 9 open reading frame 72 (*C9orf72*) is the most common genetic cause of ALS, FTD, and ALS combined with FTD (ALS-FTD) (DeJesus-Hernandez et al., 2011; Renton et al., 2011).

The clinical syndromes of FTD and ALS represent the manifestations of underlying neuropathology that results in the dysfunction and death of neurons in specific neuroanatomical regions. Therefore, individuals with bvFTD manifest with dysexecutive and behavioral symptoms because neurons within specific regions of the brain underlying executive function and social behavior (e.g., anterior cingulate, anterior insula, striatum, and amygdala) are impacted (Perry et al., 2017; Vatsavayai et al., 2019). Likewise, individuals with ALS display muscle weakness and wasting because of dysfunction and death to upper and lower motor neurons. *C9orf72*-FTD typically manifests as bvFTD (Vatsavayai et al., 2019). Anatomically, in *C9orf72*-bvFTD the cortico-striato-thalamic network (Lee et al., 2014), and medial pulvinar thalamus, specifically, appear to be the primary structures affected (Bonham et al., 2023; Sha et al., 2012; Vatsavayai et al., 2016; Yokoyama et al., 2014). Understanding the genetic landscape of normal *C9orf72* - that is, genes that are normally co-expressed with non-expanded *C9orf72* - may clarify why certain brain regions are selectively targeted in ALS or FTD (ALS/FTD), why some patients may be more likely to develop either ALS/FTD, or both; and which cell-type populations and biological mechanisms are involved.

There is considerable genetic overlap between ALS and FTD. Beyond *C9orf72,* pathogenic variants in *TARDBP, SQSTM1, VCP, FUS, TBK1, CHCHD10, and UBQLN2* (Abramzon et al., 2020) are also closely associated with both diseases. Notably, ALS, FTD and ALS-FTD patients carrying pathogenic HRE in *C9orf72* sometimes carry a second gene mutation previously implicated in ALS or FTD (Gijselinck et al., 2018). Multiple gene abnormalities in *C9orf72* HRE carriers have been detected in *TARDBP* (Chio et al., 2012; Cooper-Knock et al., 2012; van Blitterswijk et al., 2012), *TBK1* (Van Mossevelde et al., 2016), *FUS* (Millecamps et al., 2012; van Blitterswijk et al., 2012), *SOD1* (Millecamps et al., 2012; van Blitterswijk et al., 2012), *OPTN* (Cooper-Knock et al., 2012; Millecamps et al., 2012), *ANG* (Millecamps et al., 2012), *UBQLN2* (Millecamps et al., 2012), *DAO* (Millecamps et al., 2012), *GRN* (Ferrari et al., 2012), *SQSTM1* (Almeida et al., 2016), and *PSEN2* (Cooper-Knock et al., 2012; Ferrari et al., 2012; Van Mossevelde et al., 2016). Thus, the possibility of carrying a *C9orf72* HRE and a second ALS/FTD pathogenic variant is likely not random. Specific ALS/FTD genes may be co-expressed and form a functional network in brain regions that are selectively vulnerable to ALS and FTD. Consequently, disruption of these genes, depending on the affected neuroanatomical regions, is likely to influence ALS/FTD-related disease processes. Identifying genes that show similar regional co-expression patterns as *C9orf72* may, therefore, help identify novel gene targets, pathways, and biological mechanisms that mediate selective vulnerability to ALS/FTD pathogenesis.

To explore the neuroanatomical basis of shared genetic risk in the FTD/ALS spectrum, we performed gene co-expression analysis to identify genes that show regional co-expression patterns similar to *C9orf72,* the most common shared genetic contributor to ALS/FTD. We then evaluated whether the identified *C9orf72* co-expressed genes also correlate with patterns of cortical thickness in symptomatic *C9orf72* expansion carriers. Lastly, we evaluated whether certain cell populations within the brain may be selectively vulnerable to ALS/FTD pathogenesis.

## Methods

### Gene expression in the adult human brain

We investigated co-expression and regional patterns of gene expression in the healthy brain across the cortex, subcortex, and cerebellum using microarray gene expression data from the Allen Human Brain Atlas (AHBA; www.brain-map.org), a publicly available microarray dataset widely used for exploration of gene networks in the human brain (Hawrylycz et al., 2012). The microarray data was sampled from six adult human donors (3 White, 2 African American, 1 Hispanic, aged 24-57 years) in roughly 500 tissue samples from each donor, either in the left hemisphere only (n=4) or in both hemispheres (n=2). Although the same anatomical regions were sampled in all six donors, both the exact number and position of samples varied among donors. Donors also differed in other characteristics, including cause of death, post-mortem intervals, brain pH, tissue cytoarchitectural integrity, RNA quality, and number of probes used for each gene. Given space constraints, we refer the reader to the original technical white paper for additional details regarding the dissection methods, quality control, and normalization measures taken (2818165/WholeBrainMicroarray_WhitePaper.pdf).

The AHBA data were preprocessed and mapped to parcellated brain regions using a publicly available ‘abagen’ processing pipeline (https://abagen.readthedocs.io/en/stable/). We applied the recommended default parameters outlined in the original manuscripts (Arnatkeviciute et al., 2019; Markello et al., 2021). Briefly, all available probes (Custom and Agilent) were included in the analyses. Probes that did not exceed background noise in at least 50% of all cortical and subcortical samples across all subjects were excluded. As more than one probe can be available for a single gene, the probe with the higher differential stability score was selected (default parameter). Probe selection was performed for each donor separately. The Montreal Neurological Institute (MNI) coordinates of tissue samples were updated to those generated via non-linear registration using Advanced Normalization Tools (ANTs). Tissue samples were assigned to brain regions in the provided atlas if their MNI coordinates were within 2 mm of a given parcel. To reduce potential misassignment, sample-to-region matching was constrained by hemisphere and gross anatomical divisions (cortex, subcortex/brainstem, and cerebellum). All tissue samples not assigned to a brain region in the provided atlas were discarded. We used the Desikan ‘aparcaseg’ atlas (34 nodes per hemisphere + subcortex) to map tissue samples to cortical and subcortical regions (https://surfer.nmr.mgh.harvard.edu; (Desikan et al., 2006). We used the Diedrichsen atlas to map tissue samples to cerebellar regions (https://www.diedrichsenlab.org; (Diedrichsen et al., 2009). Inter-subject variation was addressed by normalizing tissue sample expression values across genes using a robust sigmoid function. Normalized expression values were then rescaled to the unit interval. Gene expression values across genes were normalized separately for each anatomic division (cortex, subcortex, and cerebellum) also using a robust sigmoid function. Samples assigned to the same region were averaged separately for each donor and then across donors. Gene expression values for the same region and gene sampled from both hemispheres from the same donor were averaged.

After implementing the pre-processing and quality control steps outlined above (Figure 1A), gene expression values from 15,633 protein-coding genes from the six donors were included in the analyses. Gene expression values were computed for 51 brain regions (34 cortical regions, 7 subcortical regions, and 10 cerebellar regions).

**Figure 1.**
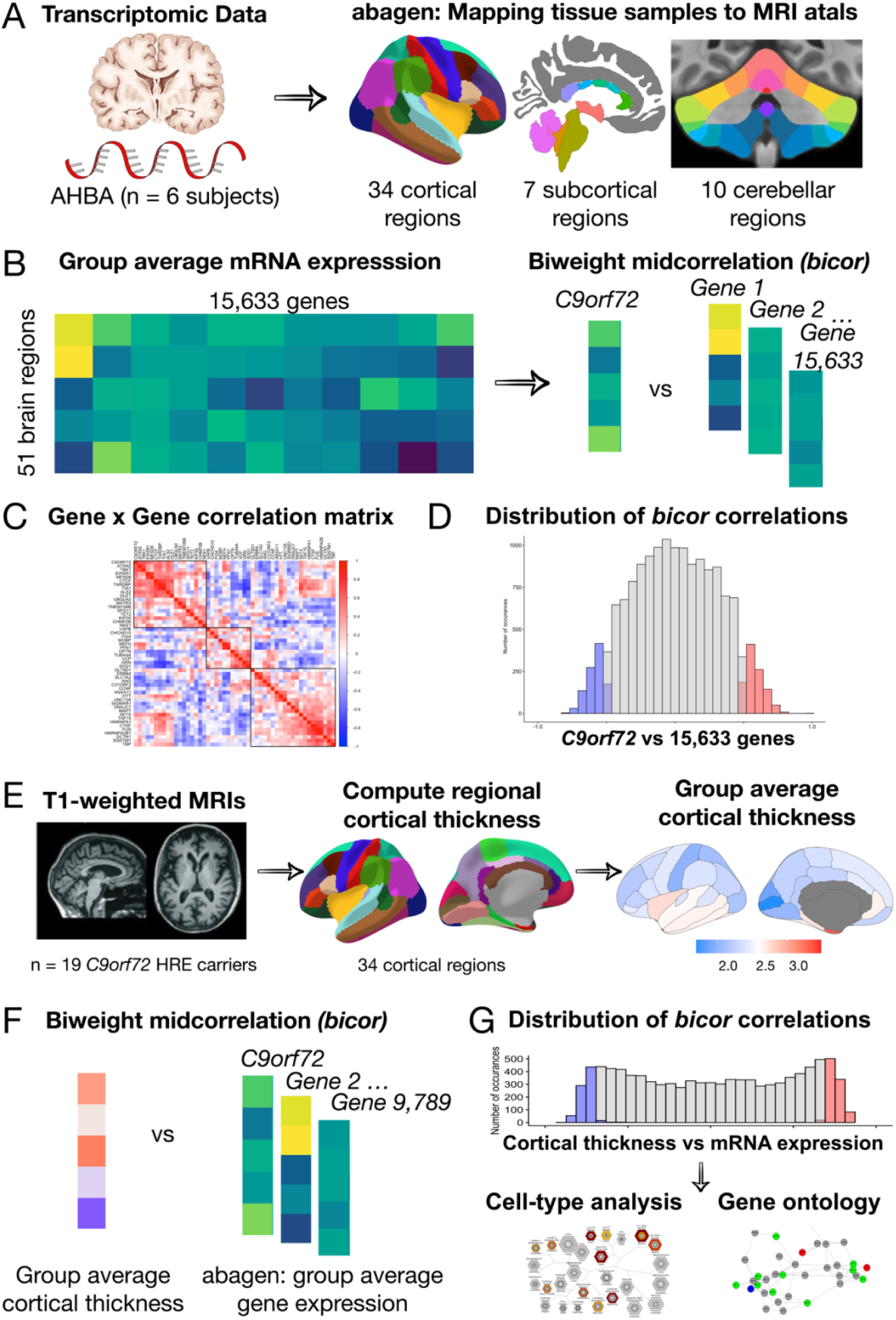
An overview of the methodology used for mapping mRNA gene expression to MRI atlas, estimation of regional gene expression, co-expression gene analysis, gene ontological and cell type analyses. **A)** Allen Human Brain Atlas samples of gene expression data were mapped to the 51 brain regions (34 cortical regions, 7 subcortical regions, and 10 cerebellar regions) according to the anatomical parcellations. **B)** Samples across the left/right hemisphere for the same region and gene were averaged across donor resulting in average gene expression values for 15,633 protein-coding genes. **C)** Using the biweight correlation method, we separately correlated the columns of each expression matrix to generate a 50 x 50 ALS/FTD gene co-expression matrix. **D)** We also used the biweight correlation method to correlate average *C9orf72* expression values for each region with average expression of all available 15,633 protein-coding genes. Subsequent analysis focused on *C9orf72*-associated genes, defined as having a correlation value above the absolute value of 0.5 (blue and red shading) (**E**) Mean cortical thickness values were extracted from 34 cortical regions for 19 patients with *C9orf72* hexanucleotide repeat expansions (*HRE*). **F)** Using the biweight correlation method, we correlated the average cortical thickness values for each region in *C9orf72* HRE carriers with average regional expression values for each *C9orf72*-associated gene. **G)** Genes that were significant at Holm-Bonferroni corrected p-value < 0.05 underwent gene ontological analyses for biological processes and cell-type enrichment analyses.

### Symptomatic *C9orf72* HRE carriers

Nineteen symptomatic *C9orf72* HRE carriers participated in this study (10 males, 9 females; age range = 48 to 81 years, mean = 64.3 years, SD = 9.7 years). Individuals were recruited from the UCSF Memory and Aging Center (MAC) FTD cohort. *C9orf72* HRE carriers were clinically diagnosed with FTD (n=8), and FTD-ALS/ALS (n=5), mild cognitive impairment (n=4), and other (n=2). For the two patients categorized as other, both showed concerns for motor neuron disease and possibly FTD. Detailed information on participant inclusion criteria can be found in prior reports (Bonham et al., 2023; Lee et al., 2014). The UCSF Committee on Human Research approved the procedures for all participants. All participants or their surrogates provided informed consent prior to participation.

### Mutation screening and genotyping

Genomic DNA was extracted from whole blood according to standard procedures. Participants were identified as carrying a pathogenic HRE in *C9orf72* if they harbored >30 hexanucleotide repeats (DeJesus-Hernandez et al., 2011). Participants in this study were negative for pathogenic variants in *MAPT* and *GRN*.

### MRI processing

All MRI scans were acquired on a 3T MRI scanner at the Neuroscience Imaging Center at UC San Francisco using previously described sequences (Bettcher et al., 2012). A high resolution T1-weighted image was acquired for all participants for structural reference, for the purpose of normalization, and for deriving morphological measures. MRI scans were processed using the FreeSurfer software package, version 6.0 (http://surfer.nmr.mgh.harvard.edu/) (Fischl, 2012). All images were visually inspected for segmentation accuracy and corrected as needed. We quantified disease burden using morphological measures of cortical thickness, since it provides a more sensitive measure of atrophy than gray matter volume (Broce et al., 2023; Winkler et al., 2010). For visualization purposes, cortical brain images were generated using the R software statistical package “ggseg,” which allows for plotting brain atlases using simple features.

## Statistical Analyses

### Statistical analysis of gene expression data

The R statistical package (version 4.2.2) was used for all statistical analysis. We first evaluated how average *C9orf72* expression in each brain region varied across different anatomical divisions: cortex, subcortex, and the cerebellum. One sample t-tests (two-tailed) were conducted to assess which of the 51 brain regions from the six donor samples expressed mRNA to a significantly greater or lesser degree compared to average mRNA expression across the whole brain. To correct for multiple tests, reported p-values were Holm–Bonferroni adjusted. Cohen’s d values for one-sample t-tests were calculated to yield a measure of effect size.

We then evaluated *C9orf72* co-expression patterns by correlating average *C9orf72* expression values for each region with average expression values from 50 other genes known to cause ALS, FTD, or combined ALS-FTD (Table 1). These genes were selected based on prior reports (Abramzon et al., 2020; Kirola et al., 2022). Two ALS/FTD genes, *PRPH* (encoding Peripherin), and *DAO* (encoding D-amino-acid oxidase), were excluded from analyses at the preprocessing stage due to low quality or coverage. We separately correlated the columns of each expression matrix to generate a 50 x 50 gene co-expression matrix, reflecting the relative expression patterns across cortical, subcortical, and cerebellar regions for each gene pair (Figures 1B-1C). We used the biweight midcorrelation as the similarity measure. In biweight midcorrelation gene expression, values that are much higher or much lower than the median value are given less weight in the correlation calculation, which makes the method more robust to extreme values and outliers (Langfelder & Horvath, 2008). Clusters were identified to assess co-expression patterns using the complete linkage method (Murtagh & Contreras, 2017).

**Table 1.**
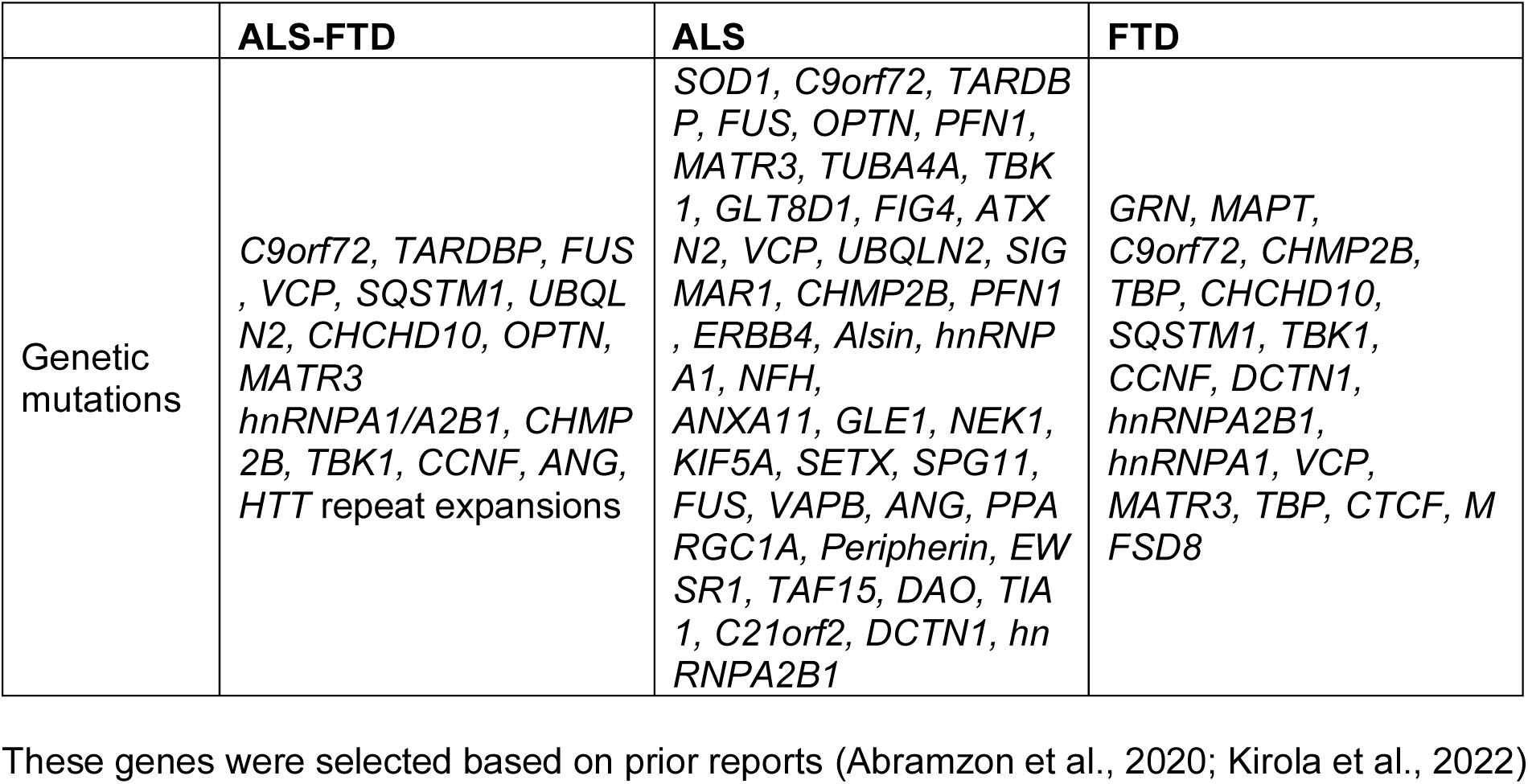

Subsequently, to screen in an unbiased manner for novel genes that are co-expressed with *C9orf72,* we used the identical procedure described above. We correlated average *C9orf72* expression values for each region with average expression of all available 15,633 protein-coding genes (Figure 1B-1D). To explore whether *C9orf72* co-expression patterns were driven by gene expression differences among the different anatomical divisions (cortex, subcortex, and cerebellum), we repeated the correlation analyses, separately, for each anatomical division. To reduce the influence of non-informative genes on subsequent analyses, genes with correlation values lower than an absolute value of 0.5 were excluded.

### Pathway enrichment analysis

We performed pathway enrichment analysis using the R statistical package ‘pathfindR’ (Ulgen et al., 2019). Since enrichment analysis of only a list of significant genes alone may not be informative enough to explain underlying disease mechanisms, we used ‘pathfindR’, which leverages interaction information from a protein-protein interaction network (PIN) to identify distinct active subnetworks and then perform enrichment analyses on these subnetworks. We preformed pathway enrichment analysis for three gene sets: “KEGG”, “GO-BP”, and “GO-MF” (all for *Homo sapiens*). For visualization, we plotted the top 20 terms for each gene set based on p-value.

## Results

### *C9orf72* gene expression patterns in the brain

Figure 2 displays how regional *C9orf72* expression varies across the cortex, subcortex, and the cerebellum. Compared to average expression across the whole brain, expression of *C9orf72* was significantly higher in seven different brain regions: pericalcarine (p-value = 0.003), VIIIb (p-value = 0.006), lingual (p-value = 0.018), inferior and superior parietal (p-value < 0.024), brainstem (p-value 0.025), and Crus II (p-value = 0.036). Expression of *C9orf72* was significantly lower in seven different brain regions: caudate (p-value = 0.0001), entorhinal cortex (p-value = 0.002), amygdala (p-value = 0.009), hippocampus (p-value = 0.014), putamen (p-value = 0.015), temporal pole (p-value = 0.016), and caudal anterior cingulate (p-value = 0.048). After Holm–Bonferroni correction, the caudate remained significant (p-adjusted = 0.006). The full results can be found in Supplementary Table 1.

**Figure 2.**
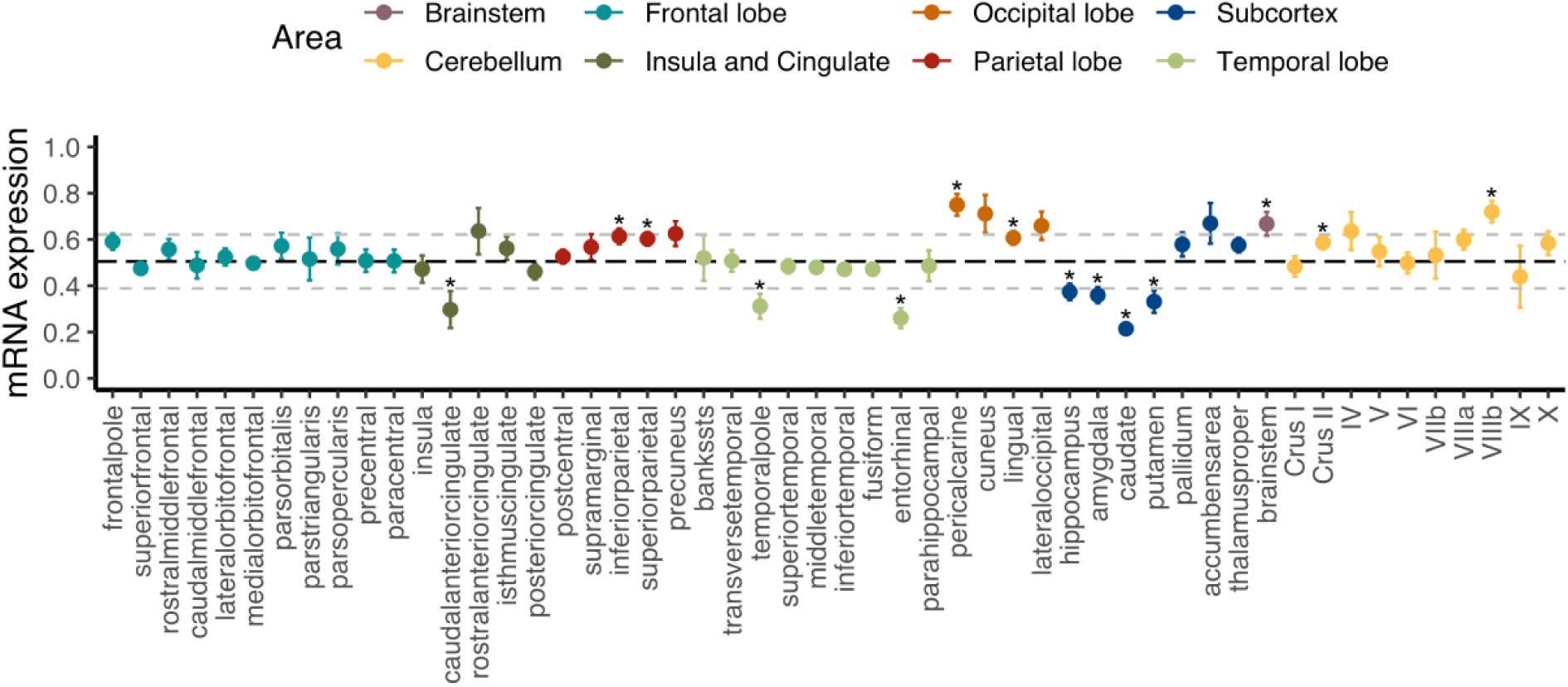
Regional *C9orf72* gene expression in the human brain. Each point represents mean expression from six donors with standard errors for a given brain region. Black bolded dashed line represents the mean expression across all genes with 1 standard deviation (+/−) also shown in light gray dashed lines. * Unadjusted *p* < 0.05

### *C9orf72* co-expression patterns in the brain

To explore whether the patterns of *C9orf72* mRNA expression in brain resemble the patterns of mRNA expression of other well-established genes known to cause ALS, FTD, or combined ALS-FTD, we correlated the average *C9orf72* mRNA expression values across the whole brain (51 brain regions) with average mRNA expression values across the whole brain for 50 ALS/FTD associated genes (Table 1). As shown in the dendrogram in Figure 3, we identified three main clusters: an orange cluster with 18 gene members, a green cluster with 11 gene members, and a blue cluster with 21 gene members. *C9orf72* was in the orange cluster and grouped with 7 other gene members: *EWSR1, MFSD8, CTCF, TARDBP, TIA, ALS2,* and *GLE1. C9orf72* mRNA expression across the brain most strongly correlated with *ATXN2* (bicor = 0.67) and *TBK1* (bicor = 0.66) mRNA expression. Interestingly, as shown in the 50 x 50 gene co-expression matrix, the orange cluster was anti-correlated with several gene members from the green and blue clusters, including *GRN, OPTN, SOD1, ANG, CCNF, VCP, SLC1A2,* among others. Overall, the strongest positive correlations in the 50 x 50 gene correlation matrix were between *HNRNPA2B1* and *FUS* (bicor = 0.86) and between *TARDBP* and *TIA* (bicor = 0.83). The strongest negative correlations were between *GRN* and *TIA* (bicor = -0.75), *SOD1* and *TIA* (bicor = -0.74), and *TARDBP* and *GRN* (bicor = -0.74). The full results can be found in Supplementary Table 2.

**Figure 3.**
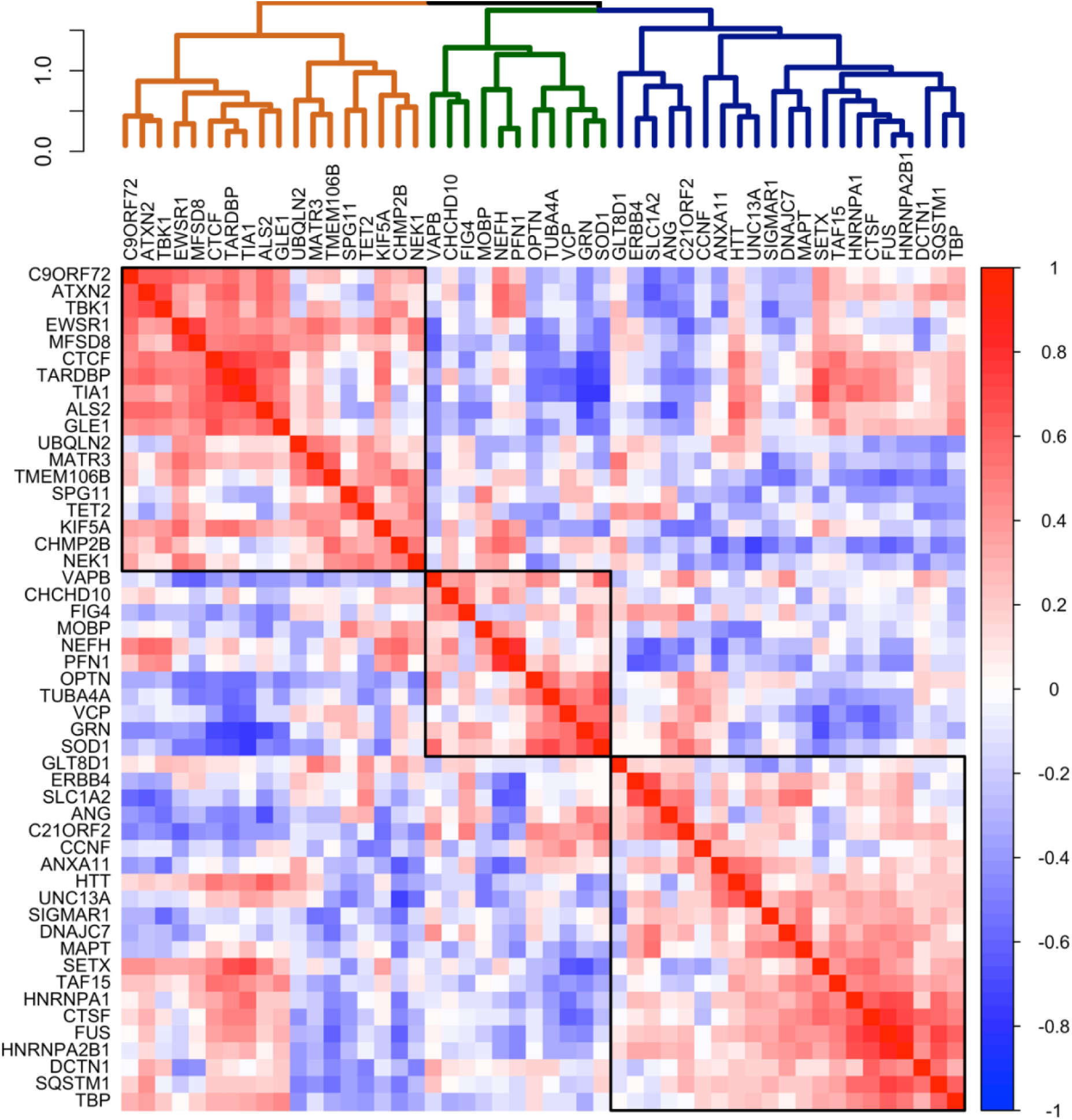
Co-expression of selected 50 ALS-FTD genes in the brain. Co-expression patterning for the expression of selected 50 ALS-FTD genes. The complete linkage method was used to identify 3 clustering groups (black squares).

### *C9orf72* co-expression patterns across different anatomical divisions

We used the same data driven approach described above to screen for new genes that were co-expressed with *C9orf72*, beyond the 50 known ALS/FTD genes. Thus, we correlated the average *C9orf72* mRNA expression values across the whole brain with average mRNA expression values for each of the 15,633 protein-coding genes. Then, to determine whether the whole brain co-expression patterns were driven by gene expression differences between the anatomical subdivisions, we visualized the similarities in whole brain co-expression and expression patterns among the different anatomical subdivisions (Figure 4A-4D). Visualizing the correlation values between *C9orf72* and a few gene members from the orange, green, and blue clusters (Figure 2) revealed that patterns across the whole brain most closely resemble the same relationships within cortical areas. Thus, for subsequent analyses, we focus on ‘*C9orf72-*co-expressed genes,’ defined by genes with a correlation value higher than the absolute value of 0.5 across one or more anatomical subdivisions (n = 9,789 genes, Supplementary Table 3).

**Figure 4.**
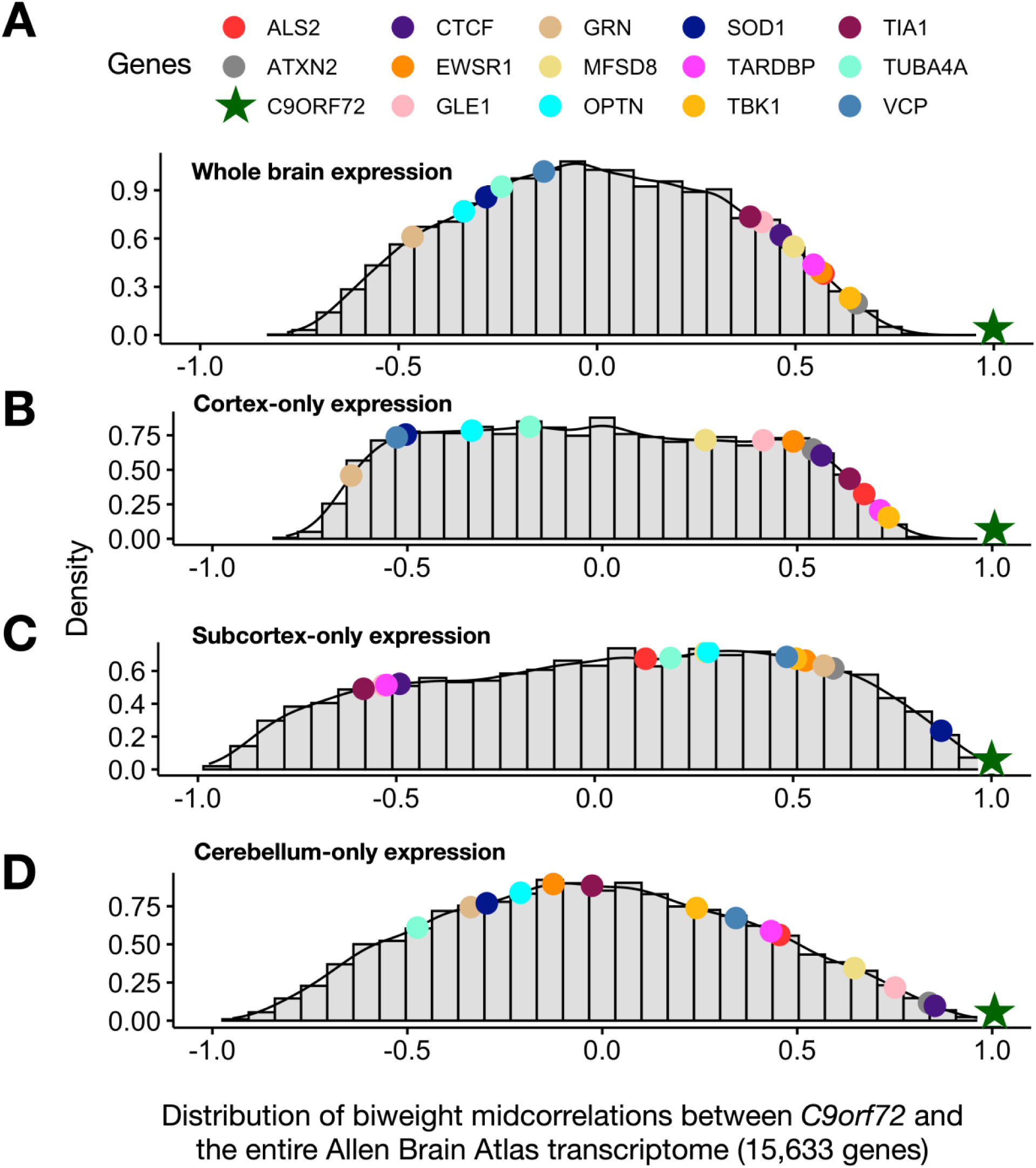
Distribution of biweight midcorrelations between *C9orf72* and all genes in the Allen Human Brain Atlas across different anatomical distributions. **A)** whole brain **B)** cortex-only **C)** subcortex-only, and **D)** cerebellum-only. Biweight midcorrelations correlations are visualized on a density distribution.

### Correlation between *C9orf72* co-expressed genes and cortical thickness in symptomatic *C9orf72* HRE carriers

A main goal of this study was to identify a network of genes or gene products that may contribute to the regional vulnerability of the human brain to *C9orf72* HRE*-*mediated pathology and, more generally, risk for ALS and FTD. Therefore, we performed correlations between average mRNA expression for each of the *C9orf72-*co-expressed genes from AHBA and average cortical thickness data from 19 symptomatic *C9orf72* HRE carriers (Figure 1E-G). Both brain mRNA expression values from the AHBA dataset and brain imaging measures from the *C9orf72* HRE carriers were mapped to the FreeSurfer average cortical surface, allowing for these imaging-genetic correlations.

After strict Holm–Bonferroni adjustment (p-adjusted < 0.05), average mRNA expression from roughly 18% of all *C9orf72-*co-expressed genes (n = 1,748 genes) significantly correlated with average cortical thickness in symptomatic *C9orf72* HRE carriers, including *C9orf72* and 15 out of the 50 ALS/FTD genes (Figure 5). Other notable correlations within the top 10 most significant include *PLEKHH3, RGS14, LRCH1,* and *PCDHB10* for their prior implication in dementia and aging (Supplementary Table 4).

**Figure 5.**
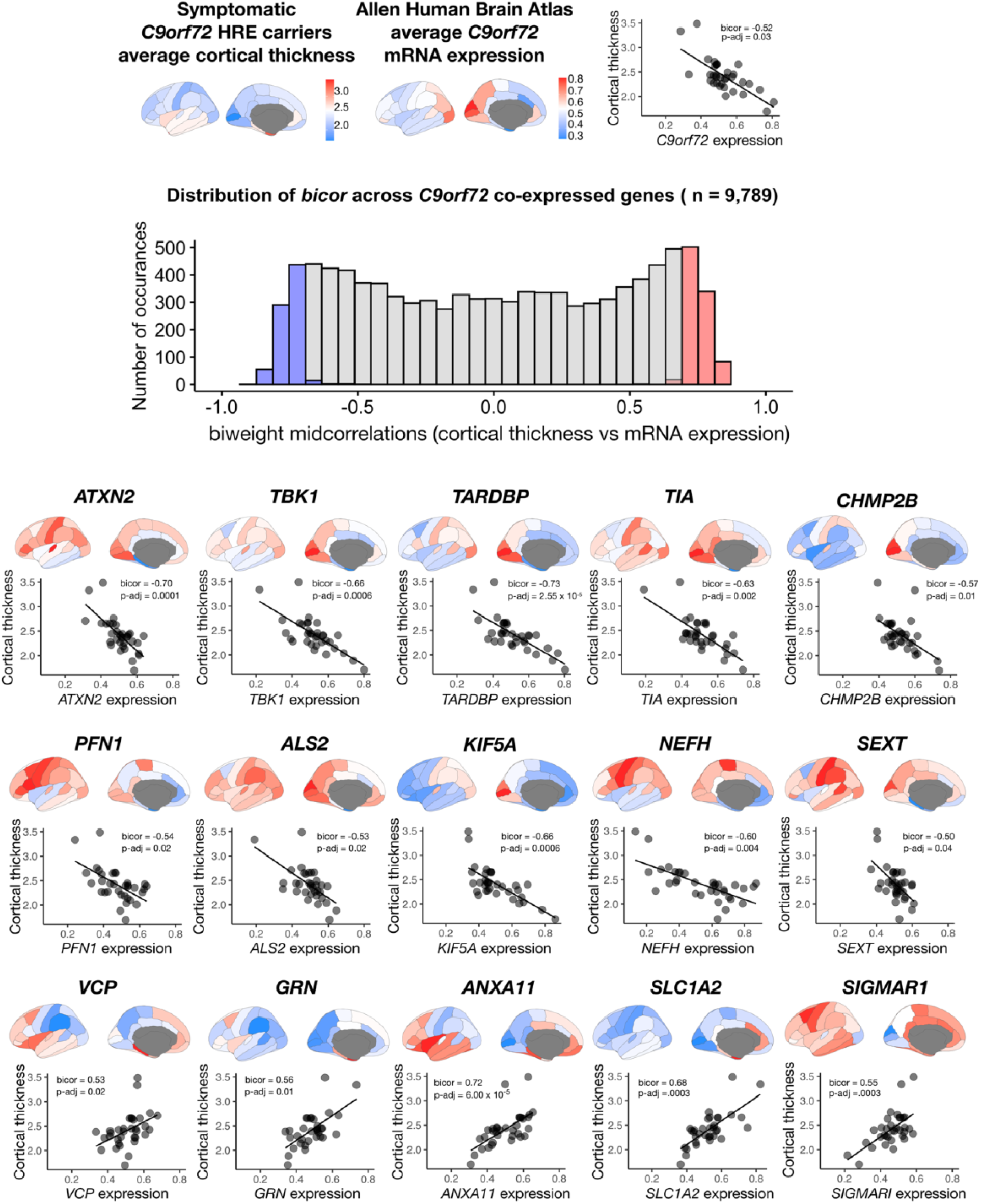
Correlation between *C9orf72* co-expressed genes and cortical thickness abnormalities in symptomatic *C9orf72* HRE carriers. **A)** average cortical thickness in symptomatic *C9orf72* HRE carriers, average *C9orf72* mRNA expression from Allen Human Brain Atlas in the same regions, and correlation between the two (top panel). **B)** After strict Holm– Bonferroni adjustment (p < .05), average mRNA expression from 1,748 genes (red and blue shading) of all *C9orf72-*co-expressed genes (n = 9,789) significantly correlated with average cortical thickness in symptomatic *C9orf72* HRE carriers. C) In addition to *C9orf72*, average mRNA expression in 15 out of 50 ALS/FTD genes significantly correlated with average cortical thickness in symptomatic C9orf72 HRE carriers.

### Evaluation of cell populations within the brain

We explored whether genes with significant *C9orf72* radiogenomic correlations (henceforth, “*C9orf72*-associated genes”) from the previous analyses were enriched for different cell populations in the brain. To do this, we used the publicly available Cell-type Specific Expression Analysis (CSEA) tools, which extracts data from the Brainspan database (http://genetics.wustl.edu/). Of the 1,748 genes we input, 1,257 were represented in the brain cell type expression dataset. These genes were enriched for several cell types or systems previously implicated in ALS and FTD (Figure 6), including layer 5b cells (Benjamini-Hochberg (BH) adjusted p-value = 0.003) (Genc et al., 2017; Nana et al., 2019; Vatsavayai et al., 2019), cholinergic motor neurons in the spinal cord (BH adjusted p-value = 0.002) (Casas et al., 2013), and Dopamine type 1 receptors (Drd1+) medium spiny neurons of striatum (BH adjusted p-value = 6.270 x 10^-4^) (Riku et al., 2016; Sobue et al., 2018), and layer 5a and immune cells (BH adjusted p-value = 0.006). Noteworthy, CSEA aimed to target Layer 5a corticostriatal interneurons, but gene expression analysis revealed substantial contamination with lymphoid cell data. For this reason, these cell lines are lumped together.

**Figure 6.**
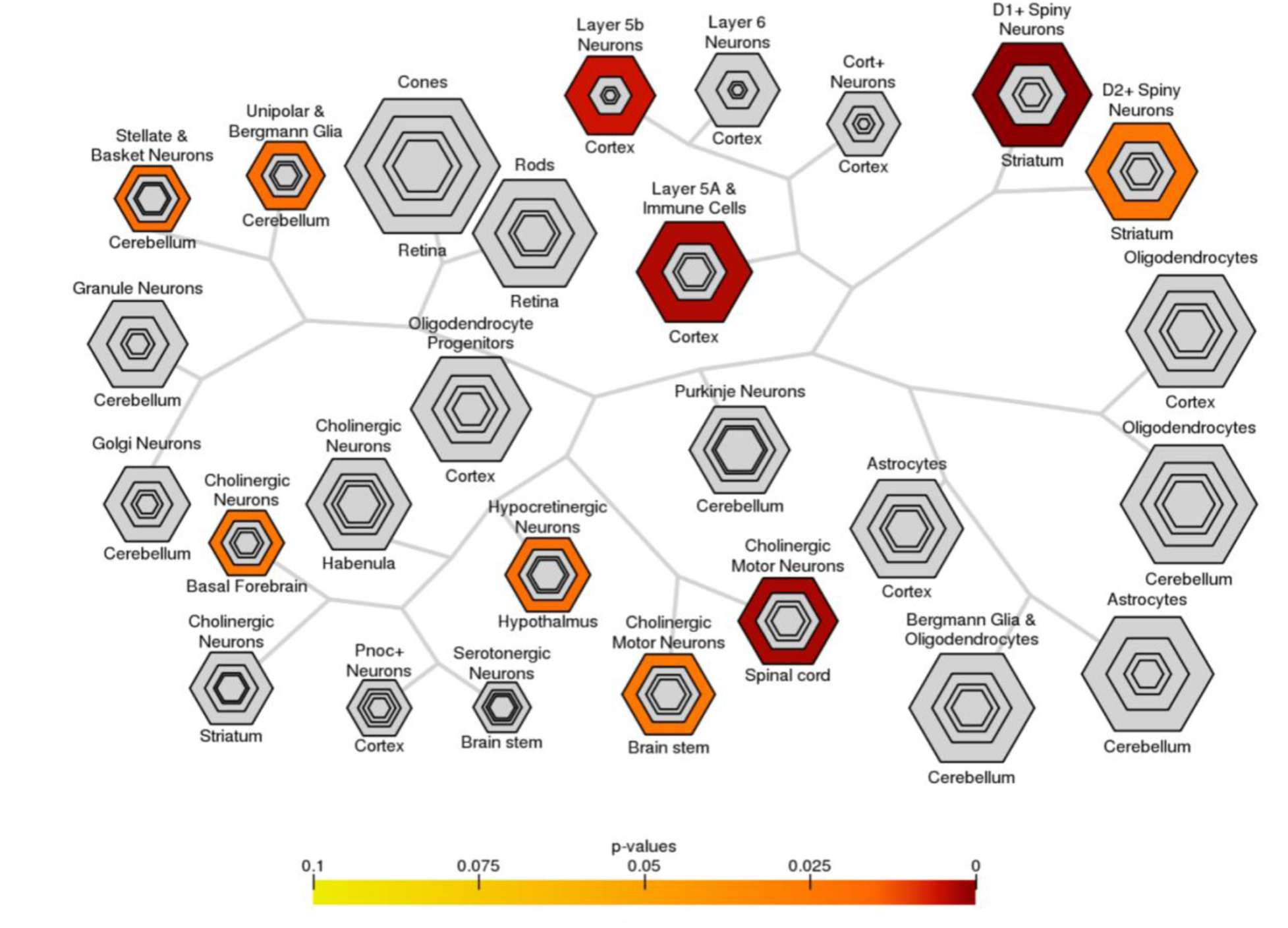
Bulls eye plot showing specific expression analysis across cell types (CSEA) of *C9orf72*-associated genes. Output of CSEA reveals a substantial over-representation at specificity index threshold (pSI) level (p-values 0.05) for layer 5b neurons, layer A and immune cells neurons, cholinergic motor neurons, and striatal medium spiny neurons. Bonferroni-Hochberg values are plotted by color.

### Identification of *C9orf72-*associated enriched KEGG pathways and GO terms

We performed pathway enrichment analysis using the R statistical package ‘pathfindR’ (Ulgen et al., 2019) to identify the biological pathways and mechanisms that *C9orf72*-associated genes were enriched for (Figure 7, Supplementary Table 5). The most relevant KEGG biological pathways included multiple synapse-related neuronal systems (glutamatergic synapse, cholinergic synapse, dopaminergic synapse, and GABAergic synapse), morphine addiction, MAPK signaling, chemokine signaling, autophagy, circadian entrainment, and mTOR signaling. Specifically, many guanine nucleotide-binding proteins (*GNAS, GNB2, GNG2, GNG4, GNG8, GNG10, GNG12*) and mitogen-activated protein kinases (*MAPK1/MAPK3*) were driving all synapse-related neuronal systems, morphine addiction, MAPK signaling, and chemokine signaling. Further, the most relevant GO biological pathway terms included autophagy and protein ubiquitination, NF-kappaB signaling, and cellular response to DNA damage. Between KEGG pathways and GO biological pathway terms, several genes were driving the autophagy enrichment: *C9orf72, TBK1, CHMP2B, VCP, MAPK1, MAPK3, C19orf12, CHMP1A, MAP1LC3B, MAP1LC3A, IRS1, PIK3CA, PIK3R4, MRAS, AKT1S1, ATG13, ATG2B, RRAGB, RRAGD, PRKCD, SH3GLB1, CAMKK2, ITPR1, ATG16L1, VAMP8 SNAP29, ULK3.* In addition to autophagy, *C9orf72* was also involved in stress granule assembly along with *ATXN2, TIA1* and late endosome to lysosome transport along with *CHMP1A* and *CHMP2B*. Lastly, the most relevant GO molecular function terms included DNA binding transcription factor activity, RNA and actin binding, and protein ubiquitination. The full list of pathways and corresponding genes can be found in Supplementary Tables 5.

**Figure 7.**
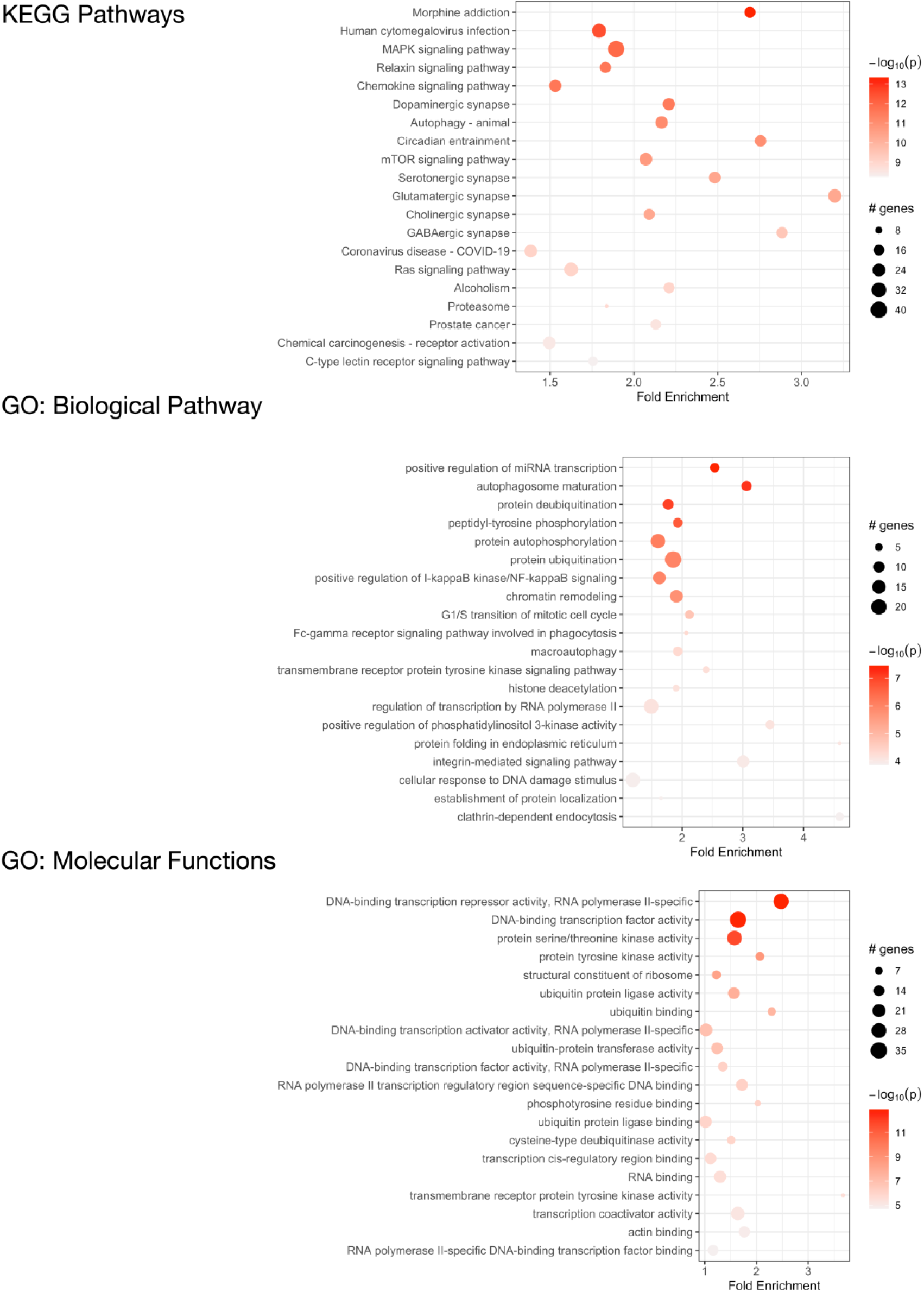
Functional enrichment analysis of *C9orf72*-associated genes. KEGG pathways that are significantly enriched in *C9orf72*-associated genes (top) and Gene Ontology (GO) analysis of biological processes (middle) and molecular functions (bottom).

## Discussion

We leveraged mRNA expression data from the AHBA and brain imaging data from 19 *C9orf72* HRE carriers to identify a network of genes and gene products that might contribute to the regional vulnerability of human brain to *C9orf72-*mediated pathology and, more broadly, to ALS and FTD. A total of 1,748 genes showed similar anatomical distribution of gene expression in the brain as *C9orf72* and significantly correlated with patterns of cortical thickness in *C9orf72* HRE carriers. These genes were enriched in expression in cell populations previously implicated in ALS and FTD, including layer 5b cells, cholinergic motor neurons in the spinal cord, and spiny medium neurons of the striatum. Despite there not being a one-to-one correspondence between gene expression levels and resultant protein levels in patients with different neurodegenerative conditions, we began to probe the potential functional relevance of the *C9orf72*-associated gene network by leveraging known protein-protein interaction networks to identify enriched pathways. The *C9orf72*-associated gene network was enriched for biological and molecular pathways associated with multiple neurotransmitter systems, protein ubiquitination, autophagy, and MAPK signaling, among others. Considered together, we identified a network of *C9orf72*-associated genes that may influence selective regional and cell-type-specific vulnerabilities in ALS/FTD.

Different brain regions have distinct distributions of cell types that are segregated into layers and have shared and unique gene expression profiles. Ultimately, these cell types form functional circuits that support motor and cognitive function, language, and social behavior. The identified 1,748 *C9orf72-*associated genes from our analyses showed selective expression in cell types previously implicated in the ALS/FTD-spectrum, including layer 5b cells in the cortex, cholinergic motor neurons in the spinal cord, and spiny medium neurons of the striatum (Figure 6). Betz cells, which are upper motor neurons, are found in layer 5b of the motor cortex. They are the largest cells in the neocortex and support long-range cortico-motor neuronal projections by sending their axons down to the spinal cord via the corticospinal tract, where they synapse directly with cholinergic lower motor neurons of the anterior horn of the spinal cord, which in turn synapse directly with their target muscle (Genc et al., 2017; McColgan et al., 2020). Damage to this system and degeneration of Betz cells and cholinergic lower motor neurons, are pathological hallmarks of ALS (Casas et al., 2013; Cleveland & Rothstein, 2001). Also located in layer 5b, but in anterior cingulate and fronto-insular cortices, are Von Economo neurons (VENs) and fork cells (Nana et al., 2019; Vatsavayai et al., 2019). VENs and fork cells are selectively vulnerable to bvFTD. The anterior cingulate and fronto-insular cortices are key regions that support social-emotional functions (Lin et al., 2019; Seeley et al., 2007; Uddin, 2015). These regions are the earliest and most consistently affected in patients with sporadic bvFTD (Kril & Halliday, 2004; Seeley et al., 2008). Also, within layer 5b of anterior cingulate and frontoinsular cortices in brain tissue of FTD patients, VEN and fork cells show disproportionate tau aggregation in FTLD-tau (Lin et al., 2019) and TDP-43 aggregation in FTLD-TDP (Nana et al., 2019; Vatsavayai et al., 2019), suggesting that VEN and fork cell biology are key aspects of FTD pathobiology. Furthermore, the *C9orf72-* associated genes were selectively expressed in medium spiny neurons of striatum. Atrophy in striatum, including caudate and nucleus accumbens, is another key feature in the ALS/FTD patients with behavioral and cognitive abnormalities, with and without *C9orf72* mutations (Masuda et al., 2016; Mirzaeva et al., 2016; Sobue et al., 2018). Corroborating these findings, previous studies have found that patients with ALS/FTD show markedly reduced striatal medium spiny neurons, particularly in the caudate head (Riku et al., 2016, 2017). In line with this work, the present study found that the caudate was the most significant region for *C9orf72* expression - *C9orf72* expression was lower here (Figure 2). Taken together, we identified a set of *C9orf72-* associated genes with selective expression in cell types known to be affected in ALS/FTD. Altering the normal gene expression profile of genes comprising the *C9orf72-*associated network may, in turn, alter the corresponding functional circuits they participate in. Thus, disruption to one or more of these functional circuits may lead to converging ALS/FTD phenotypes via different paths. Further, given that the caudate was the most significant region for *C9orf72* expression and *C9orf72* expression was lowest, *C9orf72* expression in this region may impact the intensity of *C9orf72*-specific pathological processes.

All brain cell types produce and release different kinds of neurotransmitters or chemokines which are an integral part of cell–cell communication. A subset of the *C9orf72-*associated genes, including those encoding guanine nucleotide-binding proteins (*GNAS, GNB2, GNG2, GNG4, GNG8, GNG10, GNG12*) and mitogen-activated protein kinases (*MAPK1/MAPK3*), were highly associated with multiple synapse-related neuronal system terms, such as: glutamatergic synapse, cholinergic synapse, dopaminergic synapse, and GABAergic synapse, morphine addiction, MAPK signaling, and chemokine signaling. A recent study which evaluated mRNA expression data using motor neurons derived from human induced pluripotent stem cells from ALS patients and healthy controls (Dash et al., 2020) reported that these neurotransmitter synapse-related systems were altered in their patient cohort. Notably, our results and those from Dash and colleagues demonstrate striking convergence even though the present study evaluated candidate molecular mechanisms in the healthy brain and related them to *C9orf72*-associated neuroanatomy. The support for guanine nucleotide-binding proteins and mitogen-activated protein kinase neurotransmitter genes in synapse-related systems, chemokine signaling, and MAPK signaling in ALS/FTD (Dash et al., 2020; de Oliveira et al., 2013; Sahana & Zhang, 2021) and cognitive decline in aging cohorts (Bonham et al., 2018) is not new. However, it may be that dysfunction in these neurotransmitter-associated genes may disrupt cell-to-cell communication between the specific cell-types/brain regions that we identified as associated with *C9orf72* and discussed above. This could be one way by which “throwing off” the network of genes whose proteins might be interacting at a high level in these specific brain regions could have significant consequences resulting in selective vulnerability to ALS/FTD.

The *C9orf72-*associated genes were also associated with GO biological pathway and molecular function terms in autophagy, NF-κB signaling, mTOR signaling, DNA binding transcription factor activity, and protein ubiquitination. These pathways have been consistently associated with ALS/FTD (Chua et al., 2022; Granatiero et al., 2021; Kallstig et al., 2021; Ohshiro et al., 2022; Serpente et al., 2015), and *C9orf72*-ALS/FTD specifically (Beckers et al., 2021; Serpente et al., 2015; Shao et al., 2020). By leveraging protein-protein interaction information, we found that *C9orf72* was associated with autophagy along with *TBK1, CHMP2B, VCP,* and roughly 25 other genes; stress granule assembly along with *ATXN2* and *TIA1*; and late endosome to lysosome transport along with *CHMP1A* and *CHMP2B*. Collectively, these findings have important implications for future research developing molecular-based clinical endpoints for clinical trials.

For example, a potential therapeutic strategy that may be applied for the treatment of ALS/FTD is the use of small molecules that can affect stress granule dynamics, including assembly, disassembly, maintenance, and clearance (Wang et al., 2020). Since *C9orf72*, *ATXN2*, and *TIA1* are involved in the enriched term stress granule assembly and show gene interactions (Supplementary Figure 1), developing a novel protein-based biomarker that incorporates *C9orf72*, *ATXN2*, and *TIA1* levels may help capture risk for ALS/FTD. Further, since *C9orf72, ATXN2, TIA1* cortical patterns of mRNA regional expression were strongly positively correlated with cortical thickness patterns in symptomatic *C9orf72* HRE carriers, it may be worthwhile to develop multimodal biomarkers based on cortical thickness measures and gene or protein expression data.

There are limitations to this study. Gene expression data was available using brain tissue data from healthy individuals and the imaging data were obtained from *C9orf72* HRE carriers. Therefore, the regional gene expression and imaging correlations described herein represent the relationship between regional gene expression levels in the healthy human brain and regional cortical thickness in symptomatic *C9orf72* HRE carriers. Future studies will be needed to assess regional gene expression and imaging data from the same participants.

## Conclusions

We explored the neuroanatomical basis of shared genetic risk in the ALS/FTD spectrum by identifying genes that show regional co-expression patterns similar to *C9orf72,* the most common genetic contributor to ALS and FTD. In doing so, we identified a *C9orf72-*associated gene network that was enriched in cell populations and regions within the brain known to be selectively vulnerable to ALS/FTD. Altering the normal gene expression patterns of these *C9orf72-* associated genes may disrupt cell-to-cell communication and protein-protein interactions between the specific cell-types/brain regions and, in turn, render these cell-types/brain regions vulnerable to ALS/FTD pathobiology. Future work will be required to clarify the impact of disease neuropathology on these neuroanatomical gene expression patterns.

## Acknowledgments

We thank the Memory and Aging Center at UCSF for continued support and the patients whose data contributed to this work. This work was supported by the UCSD Improving the Health of Aging Women and Men Fellowship (1T32AG058529), the Rainwater Charitable Foundation (P052319), the National Institutes of Health (NIH) (K01AG070376-03; 1R01AG058233), and the Department of Defense ALS Research Program (HT94252310353).

## Author Contributions

IJB, DWS, and JSY contributed to the concept and design of the study, analysis, and drafting a significant portion of the manuscript and figures. VES, LSS, RSD, and BM contributed to the concept and design of the study. RMN contributed to analysis of the data and generation of figures, LWB and PC contributed to the analysis.

